# Diversification and divergence in *Myioborus* warblers: insights into evolutionary relationships and plumage genetics

**DOI:** 10.1101/2025.08.12.669906

**Authors:** Laura N. Céspedes Arias, Kevin F. P. Bennett, Leonardo Campagna, Andrew W. Wood, Elisa Bonaccorso, Andrés M. Cuervo, Carlos Daniel Cadena, Irby J. Lovette, David P. L. Toews

**Affiliations:** Committee on Evolutionary Biology, The University of Chicago, Chicago, IL, USA; Birds Division, Field Museum of Natural History, Chicago, IL, USA; Department of Biology, The Pennsylvania State University, University Park, PA, USA; Department of Vertebrate Zoology, National Museum of Natural History, Smithsonian Institution, Washington, DC; Fuller Evolutionary Biology Program, Cornell Lab of Ornithology, 159 Sapsucker Woods Road, Ithaca, NY 14850, USA; Department of Ecology and Evolutionary Biology, Cornell University, 215 Tower Road, Ithaca, NY 14853, US; Natural Resources Research Institute, University of Minnesota Duluth, Duluth, MN, USA; Colegio de Ciencias Biológicas y Ambientales, Universidad San Francisco de Quito, Quito, Ecuador; Instituto de Ciencias Naturales, Facultad de Ciencias, Universidad Nacional de Colombia, Bogotá, Colombia; Departamento de Ciencias Biológicas, Universidad de los Andes, Bogotá, Colombia

**Keywords:** *evolutionary genomics*, *Neotropical mountains*, *mitogenomes*, *Ultra Conserved Elements*, *geographic variation*, *hybrid zone*, *plumage color*, *candidate gene*, *chromosomal inversion*, *CCDC91*

## Abstract

Genomic data can provide valuable insights into the evolutionary history of rapidly diversifying groups and the genetic basis of phenotypic differences among lineages. We used whole-genome sequencing of the warbler genus *Myioborus* to investigate dynamics of its recent diversification in Neotropical mountains. We found that mitochondrial and UCE phylogenies are mostly, but not fully, concordant, and we found phylogenetic support for a pattern of north-to-south and low-to-high elevation colonization in the genus. Within the *ornatus–melanocephalus* complex, which showed topological incongruence between our phylogenies, we found that genetic structure generally coincides with geographic variation in plumage, although three subspecies with striking plumage differences exhibit low mitochondrial divergence. The hybridizing taxa *M. o. chrysops* and *M. m. bairdi* show very shallow genomic differentiation, with marked peaks of divergence. Most of these are shared with other parulid warbler pairs, pointing to broad genomic features, like recombination rate, as the processes shaping these regions. However, other highly differentiated regions were unique to *Myioborus*, including one containing the gene *CCDC91*, which is associated with melanin-based plumage differences in several other birds. Lastly, we found higher levels of differentiation on the Z chromosome relative to autosomes, including two putative chromosomal inversions. Together, these results highlight the interplay of deep ancestral divergence, recent hybridization, and shared genomic architecture in shaping the evolution of phenotypic and genomic diversity within *Myioborus*.

## Introduction

Genetic data can help infer the evolutionary history of populations and species, and consequently, shed light on the processes that generate biodiversity. Since the development of multi-locus phylogenetics, it has become apparent that different genomic regions often have different genealogies (Degnan and Rosenberg 2009). This can lead to challenges when trying to establish the most plausible historical relationship among groups (Zhao et al. 2023), but also highlights the complexities of biological diversification. Multiple processes can lead to genealogical differences among loci, such as incomplete lineage sorting, selection, and introgressive hybridization, which are now appreciated as widespread and frequent phenomena across the tree of life (Rheindt and Edwards 2011; Taylor and Larson 2019). The unprecedented access we now have to whole-genome data for non-model species has allowed us to study evolutionary radiations across the tree of life in a way that considers the complex interplay between divergence and admixture (Stryjewski and Sorenson 2017; Edelman et al. 2019; Svardal et al. 2020).

In addition to reconstructing evolutionary relationships, genomic data also allow novel insights into genetic bases of phenotypic differentiation in evolutionary radiations, for example, through the description of the genome-wide landscape of divergence between phenotypically distinct taxa (Lamichhaney et al. 2015; Campagna et al. 2017; Marques et al. 2022), which are often heterogeneous (i.e. with distinct peaks and valleys). Although the causes of the heterogeneity of landscapes of divergence are multiple and complex (e.g., variation in recombination rates, variation in mutation rates; Ravinet et al. 2017), identifying divergence peaks is a useful first approach to identify candidate genomic regions related to phenotypes of interest. This approach has proven useful to study several avian species complexes and has allowed significant progress in identifying candidate genes underlying melanin and carotenoid-based plumage traits (Toews et al. 2016; Campagna et al. 2017; Aguillon et al. 2021; Baiz et al. 2021; Funk et al. 2023; Lim et al. 2024).

The parulid warblers of North, Central, and South America comprise a speciose group of colorful passerines that radiated over the last 7 MY (Rabosky and Lovette 2008; Oliveros et al. 2019). One of the most conspicuous phenotypic differences across the parulid family is plumage color, with little morphological variation (Shutler and Weatherhead 1990; Leroy et al. 2024) (but see Rosamond et al. 2020), in contrast to classic adaptive radiations characterized by marked differences in size and shape, e.g., *Anolis* lizards, Darwin’s finches, African cichlids, Hawaiian honeycreepers, plants in the genus *Espeletia* (Losos 1990; Seehausen 2006; Foster et al. 2008; Lerner et al. 2011; Tokita et al. 2017; Pouchon et al. 2018). Because evolutionary radiations produce many species quickly, the resulting lineages are usually genetically compatible, thus allowing for hybridization and gene flow (Seehausen 2004; Abbott et al. 2013). This phenomenon is true for parulid warblers (Baiz et al. 2021), where several hybrid zones have been described, and numerous hybrids have been documented (Short and Robbins 1967; Rohwer and Wood 1998; Vallender et al. 2007, 2016; Irwin et al. 2009; Toews et al. 2011, 2020; Céspedes-Arias et al. 2021).

Given their recent diversification, combined with extensive hybridization among lineages, one may expect a variety of gene genealogies across loci in warblers (Singhal et al. 2021). Thus, it is necessary to leverage whole-genome data and use phylogenomic approaches to appropriately describe this groups’ evolutionary history. In this study, we focused on a lineage of rapidly diversifying warblers (Lovette et al. 2010): the genus *Myioborus*, composed of 12 colorful, mostly montane, species occurring from the southern United States to Argentina. This genus is one of several lineages of resident warblers that have diversified in South and Central America, which as a whole make up more than half of the species diversity in Parulidae (Curson 2010; Lovette et al. 2010; Winger et al. 2012). Despite their remarkable diversity, comparatively little genomic data has been generated for these tropical resident warblers, in contrast to migratory species that breed in North America (e.g. Baiz et al. 2021; Zamudio-Beltran et al. 2024).

Our study focuses on *Myioborus* warblers to both infer the evolutionary relationships between species and geographically circumscribed plumage groups, as well as to describe the landscape of divergence in a young species pair characterized by conspicuous differentiation in color patterns. Most *Myioborus* species have allopatric distributions across mountain ranges, or parapatric ranges along elevational gradients. Ten species occur exclusively in upper montane forest (hereafter, high-elevation *Myioborus*), including representatives in Central American cordilleras, the Sierra Nevada de Santa Marta, the Tepuis, and the tropical Andes. This group is characterized by plumage coloration diversity that is concentrated in the face and crown, with species showing combinations of rufous, black, and yellow crown colors, as well as presence or absence of black, yellow, or white cheeks, lores and eye rings, forming “spectacles” (Figure 1). A previous examination of evolutionary relationships among *Myioborus,* which identified this upper montane taxa as forming a monophyletic group, was based on a few mitochondrial loci and lacked resolution inferring relationships within this clade, potentially as a result of rapid diversification (Pérez-Emán 2005). Phylogenetic studies of the entire warbler family have provided insights into the diversification of *Myioborus* taxa, albeit with some limitations due to limited geographic sampling within geographically variable species in this upper montane clade (Lovette et al. 2010; Zhao et al. 2025). The use of a comprehensive taxon and geographic sampling of genomic data within *Myioborus*, especially within this upper montane clade, is necessary to better understand how these species have diversified across Neotropical mountains.

**Figure 1.**
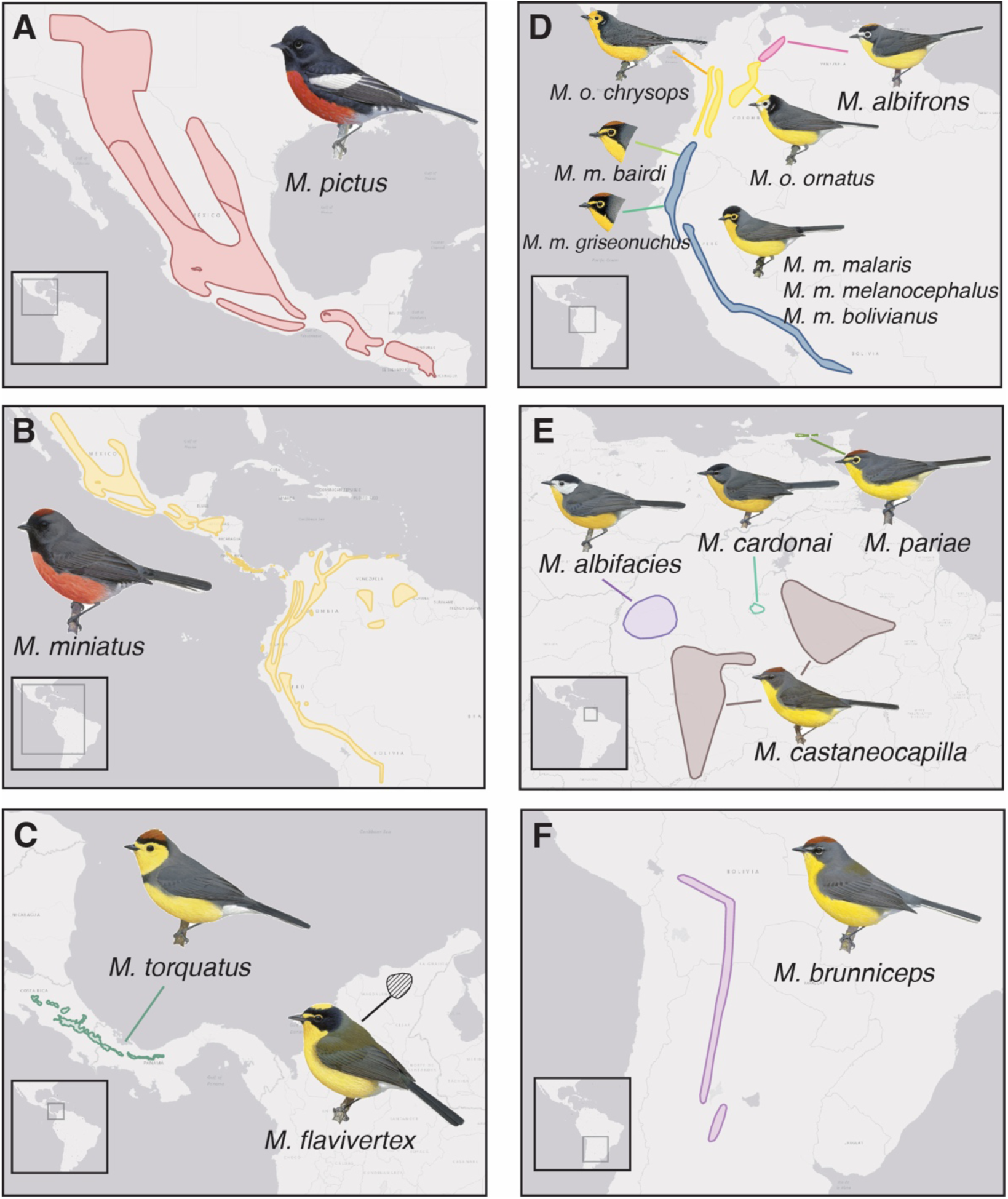
Distribution and plumage diversity in the genus *Myioborus*. Panels are focused on different geographic regions of Central and South America, with some including multiple species. Panel D corresponds to the species *M. albifrons*, and the geographically variable *M. ornatus* and *M. melanocephalus* (sampling sites for these species in Figure 3). The approximate distribution of species that we sampled is shown as colored polygons; the distribution of the only species in the genus not included in the study is shown in a hatched pattern in panel C (shapefiles from NatureServe). Warbler illustrations used with permission, © Lynx Nature Books and © Cornell Lab of Ornithology.

One species complex (*Myioborus ornatus-M. melanocephalus*) within this high-elevation group is distributed from northern Bolivia to western Venezuela and displays striking geographic patterns of plumage variation (Pérez-Emán 2005; Lovette et al. 2010). These two species comprise between five and seven well-defined plumage groups depending on taxonomic designations (Cuervo and Céspedes Arias 2023). Data from a single mitochondrial locus (Céspedes-Arias et al. 2021) suggest that the phenotypically diverse *ornatus*-*melanocephalus* group is overall characterized by low genetic differentiation, implicating either rapid recent diversification, introgression, or both. Given the low genetic differentiation within the *ornatus-melanocephalus* complex, whole genome data are needed to better understand how genetic structure correlates with the observed geographic variation in plumage color. Recent work, using comprehensive geographic sampling and reduced-representation genomic data, suggests that patterns of plumage and genetic differentiation are largely concordant. This study also indicates that both topographic barriers within the cloud forest and isolation by distance contribute to shaping genetic variation within this complex (Céspedes Arias, Campagna *et al,* preprint).

Two of the plumage groups within this species complex form an extensive hybrid zone in the Colombia-Ecuador border region, where intermediate plumage phenotypes are common and diverse (Céspedes-Arias et al. 2021). Given low divergence in previous genetic surveys (Céspedes-Arias et al. 2021, Céspedes Arias, Campagna *et al,* preprint), by using whole-genome resequencing data we can further examine wether this shallow level of divergence is also observed in the nuclear genome, and how the levels of divergence vary across the genome, potentially allowing us to identify candidate genes for their striking head plumage pattern differentiation.

Here we obtained whole-genome resequencing data representing 11 of the 12 species in *Myioborus* (excluding *M. flavivertex*, a species endemic to the Santa Marta mountains of Colombia). We inferred the evolutionary relationships among species and plumage groups and evaluated topological incongruence between the nuclear and mitochondrial genome resulting from introgression or incomplete lineage sorting. We next addressed cases of tree incongruence, both between previous studies (Lovette et al. 2010; Zhao et al. 2025) and between our tree-building methods. Additionally, we described nuclear and mitochondrial divergence among *M. ornatus*, *M. melanocephalus*, and *M. albifrons*. Finally, we focused on the youngest taxa in the genus, *M. o. chrysops* and *M. m. bairdi*, which hybridize in secondary contact, and identified genomic features of divergence between them, including a gene putatively underlying plumage color.

## Methods

### Study species and sampling

We generated short-read whole-genome resequencing data (n=57 in total, Supplementary Table 1) for 11 out of 12 species in the genus *Myioborus*: *M. pictus* (n=5), *M. miniatus* (n=5), *M. torquatus* (n=5), *M. pariae* (n=1), *M. castaneocapilla* (n=5), *M. cardonai* (n=2), *M. albifacies* (n=1), *M. brunniceps* (n=4), *M. albifrons* (n=2), *M. ornatus* (n=11), and *M. melanocephalus* (n=14). Most of the tissue samples used were loaned from museum collections in the US (AMNH, FMNH, KU, LSUMZ, MSB, UWMB, and YPM; see Supplementary Table 1 for the full names). A subset of samples (n=18) from the latter two species correspond to the hybridizing taxa *chrysops* and *bairdi* of the Tropical Andes, and come from a previous sampling effort by Cespedes *et al*. 2021 (see for more detail in collection methodology). All tissue samples used here are associated with vouchered museum specimens (Supplementary Table 1).

A higher proportion of samples correspond to the polytypic species *M. ornatus* and *M. melanocephalus*, which allowed us to have a good representation of their taxon diversity and geographic variation in plumage pattern and color. The southern species *M. melanocephalus* traditionally comprises five subspecies (from south to north; Figure 1): *bolivianus*, *melanocephalus, malaris, griseonuchus* and *bairdi* (formerly *ruficoronatus,* see Cuervo & Céspedes Arias 2023). These taxa differ in their crown color (either black or rufous) and the extent of black in the lores and forehead that delineate “spectacles” characteristic of the species (for detailed descriptions see Cuervo and Céspedes Arias 2023). The northern species *M. ornatus* lacks these “spectacles” and is instead characterized by a yellow forehead and either yellow (subspecies *chrysops*) or white cheeks (subspecies *ornatus*) (Figure 1). We obtained whole genome re-sequencing data for 6 out of 7 of these subspecies, with higher sample sizes (n=9) for *M. o. chrysops* and *M. m. bairdi*, the two taxa that hybridize extensively in the Colombia-Ecuador border. The specimens we included from this taxon pair correspond to males with clearly parental phenotypes based on the plumage scores of Céspedes-Arias et al. (2021). We sampled from across the subspecies’ ranges, including several from the north and south edges of the hybrid zone that had parental phenotypes.

### Sequencing methods

After extracting genomic DNA, we conducted low coverage re-sequencing of the genomes of 57 *Myioborus* warblers. For the 18 samples collected by Cespedes *et al* 2021, we conducted laboratory work in the Fuller Evolutionary Biology Lab (Cornell Lab of Ornithology). We prepared libraries barcoded by individual using the TruSeq Nano DNA Library Prep kit (Illumina, California, USA) with a 550 bp insert size. These samples were then pooled and sequenced in one Illumina NextSeq500 (2 x 150 bp) lane at the Cornell Institute of Biotechnology. For the remaining 39 samples, laboratory work was conducted at the Toews Lab at Pennylvania State University. Libraries were also prepared using Illumina DNA Prep kits and these samples were pooled, together with samples from other projects, and subsequently sequenced across five NextSeq P3 Illumina flowcells on a NextSeq 2000 at the Genomics Core Facility within the The Huck Institutes of Life Sciences.

### Bioinformatic processing of raw reads

First, we trimmed adapters and collapsed overlapping reads with AdapterRemoval 2.17 (Schubert et al. 2016). The number of raw reads per sample ranged from 11.2 to 47.5 million (mean= 23’773,417; SD= 6’790,475; see Supplementary Table 1 for per sample values). We then aligned the reads to the chromosome-level *Setophaga coronata* (Baiz et al. 2021) reference genome using Bowtie2 (Langmead and Salzberg 2012). The resulting SAM files were then converted to BAM and sorted using Samtools (Li et al. 2009), and PCR duplicates were marked using Picard (Broad Institute 2019). The depth of coverage of the resulting assemblies ranged from 2.4 to 8.8X (mean= 3.9; SD= 1.05; Supplemtary Table 1), and the the alignment rates ranged from 0.84 to 0.89 (mean= 0.87; SD= 0.01; Supplemtary Table 1).

### Analyses

To infer evolutionary relationships among *Myioborus* species and plumage groups, we conducted phylogenetic inference using mitochondrial genomes and nuclear loci, specifically Ultra Conserved Elements (UCEs, (Faircloth et al. 2012)). Then, to describe the genetic structure within the *ornatus-melanocephalus* complex and how it relates to plumage variation, we built haplotype networks with the mitochondrial genomes and conducted principal component analyses to summarize the genetic variation across nuclear sites. Lastly, we describe the landscape of divergence between hybridizing *chrysops* and *bairdi* by estimating F_ST_ values per genomic window, and statistically inferred outliers (i.e. high divergence peaks) and those that are unique to these *Myioborus* pair relative to other comparisons in the warbler family. Below we described how we obtained the mitochondrial genomes and nuclear genome data (UCEs and genotype-likelihoods for millions of sites) and how we conducted these analyses.

#### Mitochondrial genomes

To obtain a mtDNA VCF we first used bcftools *mpileup* (Li et al. 2009) subset to retainreads mapping only to the *S. coronata* mitochondrial sequence retrieved from the reference genome, and then used bcftools *filter* (Li et al. 2009) to set any genotype with a genotype quality (GQ) less than 10 to uncalled. We then obtained a consensus FASTA sequence for each sample, using the functions samtools *faidx* (to extract the mitochondrial genome from the reference genome) and bcftools *consensus* (to build the FASTA files using the extracted reference mitochondrial region and the VCF as input). Finally, we aligned the individual FASTA files using mafft (Katoh et al. 2002).

*mtDNA phylogenetic tree and haplotype network-* We used the mtDNA alignment to infer a phylogenetic tree with Iqtree2 (Minh et al. 2020), using ModelFinder Plus (-m MFP), and collapsing near-zero-length branches (-czb). We assessed branch support by using 1,000 replicates of an approximate likelihood-ratio test (-alrt 1000). In addition, we built the network of mitochondrial haplotypes within the *M. ornatus-M. melanocephalus* complex using the function haploNet from *pegas* (Paradis 2010). To import and manipulate the alignment in R we also used functions implemented in *ape* (Paradis and Schliep 2019)

#### Nuclear genomes

##### UCE (Ultra Conserved Element) phylogenetic trees

We inferred phylogenetic relationships among *Myioborus* species using UCEs, which have proven useful to resolve phylogenetic relationships both deep and shallow time scales (Carter et al. 2023). To conduct phylogenetic analyses, first we performed variant calling. We combined individual alignment files into a master file using using bcftools *mpileup,* and the *S. coronata* genome as reference. Then we called variants using bcftools *call,* specifying that the genotype quality should be included in the output (*-f GQ*) and that these correspond to diploid samples (*--ploidy 2*). Lastly, we left-aligned and normalized indels using bcftools *norm*. Next, using bcftools *filter,* we set the genotypes with a GQ lower than 10 as missing. This final VCF consisted of 959.2 million sites, 92.5 million SNPs and 3.7 million indels.

We used a set of 4016 UCEs and flanking sequences totaling 2 kb each in length that were previously identified in the S. coronata assembly (see Baiz et al. 2021). To extract the UCEs from the filtered VCF, we first created a VCF only including UCE sites using bcftools *view*, and then created species-fasta files containing each 2-kb region using the approach described above (using samtools *faidx* and bcftools *consensus*). Then, we aligned and filtered UCEs using phyluce (Faircloth 2016). We first aligned the UCE fasta files using *phyluce_align_seqcap_align*, specifying mafft (Katoh et al. 2002) as the aligner, and then did internal trimming with GBlocks. All 4016 UCE loci were present for all taxa, so we did not drop any. Finally, we concatenated the UCE alignments and output a single phylip file using *phyluce_align_concatenate_alignments*.

Before running Iqtree2 (Minh et al. 2020) to build the UCE tree, we determined the best molecular evolution model for our data using a subset of individuals for computational feasibility. To do this, we followed the same alignment steps described above only for 11 individuals representing all sampled *Myioborus* species and one outgroup (*Cardellina pusilla*). Then we ran Iqtree2, using the flag *-m MFP*, to enable model selection with ModelFinder Plus. Based on the result from this run, we used the TVM+F+I+R7 for the Iqtree2 run including the full set of individuals. For the full run, we specified 1,000 approximate likelihood ratio test replicates to assess branch support (*-alrt 1000*).

To visualize possible discordance between the mitchondrial and the UCE (nuclear) phylogenetic tree, we generated a cophylo plot using the *cophylo* function implemented in the *phytools* library version 2.0 (Revell 2024) for R.

In addition to the concatenated UCE tree, we also used gene tree reconciliation to make a species tree. For this method, we made trees for each UCE locus individually in iqtree using the best fit sequence evolution determined above, then used those as inputs to ASTRAL-III (Zhang et al. 2018), which builds a species tree weighted on branch length and support values of input trees.

##### Genotype likelihood-based PCAs

To further explore genomic variation in the Tropical Andean (*ornatus-melanocephalus-albifrons*) clade, where we had a better geographic sampling including all major plumage groups, we conducted a PCA in ANGSD (Korneliussen et al. 2014) (12,624,354 sites). First, we calculated genotype likelihoods using the *angsd* command and specifying the GATK likelihood model (*-GL 2*). We also specified that only SNPs with a p-value equal or lower than 1e-6 would be called (*-SNP_pval 1e-6*). Then we used the python script *pcangsd.py,* which uses as input the genotype likelihoods in BEAGLE format generated in the previous step to perform the PCA.

To generate a PCA for the *ornatus-melanocephalus* complex only, we followed the same procedure described above but excluding those individuals corresponding to *M. albifrons* (13,314,393 sites).

##### Landscape of divergence between hybridizing *Myioborus*

To describe the landscape of divergence between the hybridizing taxa *chrysops* and *bairdi* (Céspedes-Arias et al. 2021), and potentially identify genomic regions underlying plumage differences, we calculated windowed F_ST_ using the realSFS program from ANGSD. First, we generated the site frequency spectrum from allele frequency likelihoods (realSFS), and then we prepared the data for the calculation of F_ST_ (realSFS *fst index*). Finally, we calculated F_ST_ values (realSFS *fst stats2*) using non-overlapping sliding windows of 10 kbp (-win 10,000, -step 10,000).

Divergence peaks between taxa can result from multiple factors in addition to regions underlying differentiated traits. Processes that may operate deeper in evolutionary time and which generate regions of reduced recombination or low genetic diversity can produce common “landscapes of divergence” (Cruickshank and Hahn 2014). For instance, Baiz et al. (2021) studied nine independent species pairs of *Setophaga* and found 43% divergence peaks shared across two or more pairs, likely the result of divergence acting upon an ancient and shared genomic substrate. To separate divergence peaks that are likely to underlie unique trait differences between the hybridizing pair *chrysops* and *bairdi* from those that are likely attributable to other evolutionary processes operating at deeper time scales (Burri et al. 2015), we compared 10 kb windowed F_ST_ estimates from our study taxa to those from Baiz et al. (2021).

Specifically, we identified which of the top 0.1% of 10-kb *Myioborus F*_ST_ windows, calculated separately for the Z chromosome and autosomes, fell outside the 1 Mb regions which Baiz et al. (2021) determined contained an overrepresentation of elevated F_ST_ windows between one or more sister pairs of *Setophaga* warblers. We also compared each windowed F_ST_ value in *Myioborus* to the same window in the comparison between the non-hybridizing sister species *S. striata* and *S. castanea* from Baiz et al. (2021)—a pair that has moderate levels of genetic differentiation and shares many divergence peaks with other *Setophaga* warbelrs—to identify divergent regions unique to the *Myioborus* pair on a finer scale.

Because we found notable patterns of divergence between *chrysops* and *bairdi* in the Z-chromosome (see below), we further examined genetic variation in this chromosome at the individual level by conducting a region-specific PCA. To do this, we manually subsetted the genotype likelihood beagle file to four regions spanning two Z chromosome divergence peaks (12,233 and 21,398 sites) and two adjacent regions of low divergence (136,312 and 59,678 sites). Then we conducted the PCA in ANGSD, as explained above for the full genome VCF file.

## Results

### Phylogenetic relationships in *Myioborus* inferred using mitochondrial genomes and UCEs

The topologies of the mtDNA and both the UCE phylogenies (i.e. the concatenated UCE tree and the species tree inferred using ASTRAL) were mostly congruent (Figure 2, Figure S3), with some exceptions. In all three phylogenies, the earliest two diverging branches contain *M. pictus* and *M. miniatus*, respectively, with *M. miniatus* sister to all upper motane *Myioborus*. All three methods supported a deep divergence between Central and South American *M. miniatus*, but while the mtDNA tree recovered *M. miniatus* as monophyletic, the UCE trees supported South American *M. miniatus* as sister to all remaining species in the high elevation clade, all trees showing high branch support (Supplementary Figures 1 and 2).

**Figure 2.**
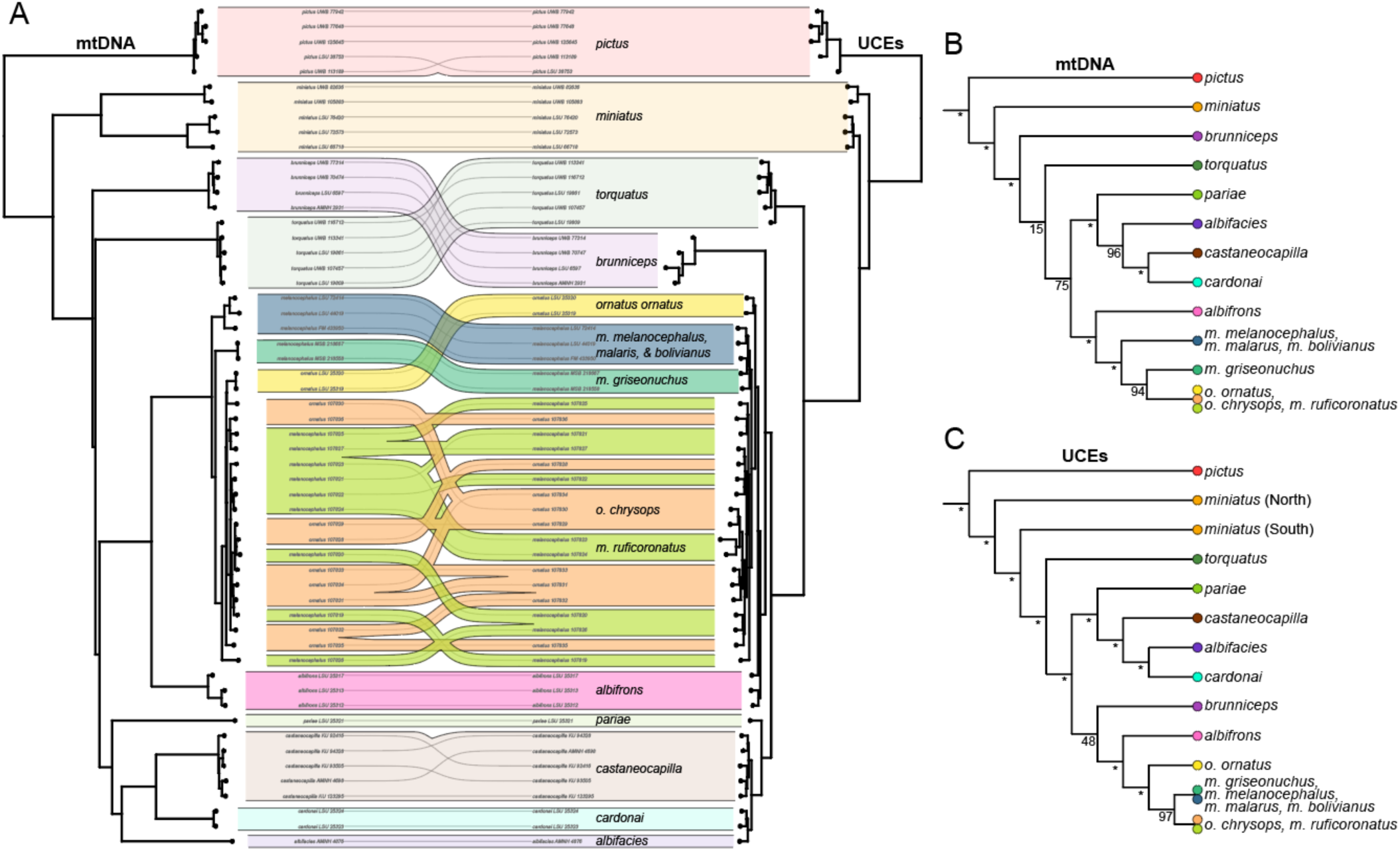
Phylogenies inferred based on the mitochondrial genome and nuclear loci (UCEs) show mostly concordant topologies. Color codes as in the remaining figures, pink= *M. albifrons*; yellow=*M. o. ornatus*; orange= *M. o. chrysops*; lime green= *M. m. bairdi;* aquamarine= *M. m. griseonuchus*; blue= *M. m. malaris*, *M. m. melanocephalus* and *M. m. bolivianus* **A)** Cophylogeny plot highlighting the topological differences between the mtDNA and concatenated UCEs phylogenies. The lines in the middle connect the same individual sample in both phylogenies. **B)** Simplified topology of the mtDNA tree. **C)** Simplified topology of the UCEs concatenated tree. In both B and C support values are shown, with the asterisk (*) representing 100.

Within the upper montane *Myioborus* group, the mtDNA and UCE trees consistently recover two well-supported clades: a tropical Andes clade containing *M. albifrons, M. ornatus,* and *M. melanocephalus* and a Tepui region and coastal ranges of Venezuela clade containing *M. pariae, M. castaneocapilla, M. cardonai,* and *M. albifacies* (Figure 2, Supplementary Figs. 1 and 2). Within this major upper montane clade, however, there is some uncertainty in the placement of *M. brunniceps* (subtropical Andes) and *M. torquatus* (Central America). The mtDNA tree positions *M. brunniceps* as sister to all other high elevation species, though with low branch support (15%, Supplementary Figure 1). In contrast, the UCE trees place *M. torquatus* in this position with strong support. The concatenated UCE tree recovers a tropical Andes plus *M. brunniceps* clade with low support (15%, Supplementary Figure 2), whereas the ASTRAL UCE tree places *M. brunniceps* as sister to the tropical Andes plus Tepui clade with strong support (100%, Figure S3).

Within the Tepui clade, all three trees placed *M. pariae* as sister to the rest of the clade. The ASTRAL UCE tree was unable to resolve more recent splits in either the Tepui or tropical Andes clades—with less than 90% support for all species relationships—possibly owing to low phylogenetic signal between recently diverged lineages. The mtDNA and concatenated UCE phylogenies differed in the sister relationship of *M. cardonai*: the mtDNA tree showed *M. cardonai* sister to *M. castaneocapilla*, while the UCE tree recovered *M. albifacies* as sister to *M. cardonai.* Similarly, the Andean group also showed some discordance between trees. In this group, for which we had the most comprehensive sampling, both trees supported *M. albifrons* (Venezuelan Andes species) as sister to a clade encompassing all *ornatus-melanocephalus* populations. This result reflects the Lovette et al. (2010) topology rather than the Zhao et al. (2025) toplogy, possibly owing to a poor-quality *M. ornatus* sample from Zhao et al. (2025). Our UCE and mtDNA phylogenies differed, however, in the placement of white-faced *M. o. ornatus* inhabiting the Eastern Andes of Colombia up to the Táchira depression in Venezuela (Figure 1). In the mtDNA tree, *M. o. ornatus* was embedded in a clade with samples of *M. o. chrysops* and *M. m. bairdi*, supporting a clade of northern *ornatus-melanocephalus* taxa ranging from Ecuador to western Venezuela. In contrast, in the concatenated UCE tree, *M. o. ornatus* appeared outside of the Colombia-Ecuador clade (*M. o. chrysops, M. m. bairdi*), as sister to all *melanocephalus-ornatus* populations, including the southern forms.

### Population genetic structure in the *ornatus-melanocephalus-albifrons* group

Where phylogenetic methods fail to clearly identify evolutionary relationships, whole-genome sequence data can help shed light on broad and fine-scale patterns of genetic divergence between taxa. We therefore focused on the Tropical Andes clade (*M. albifrons, M. ornatus, and M. melanocephalus*), which has been difficult to resolve and for which we had greater sampling.

The haplotype network of the mitochondrial genomes (Figure 3) showed a few identifiable clusters, roughly consistent with the mtDNA tree. Specifically, it shows a cluster of all *M. albifrons* individuals, a cluster of samples corresponding to *M. m. melanocephalus, M. m. malaris* and *M. m. bolivianus* (from central and south Peru, all in dark green), a cluster corresponding to *M. m. griseonuchus* (from northern Peru, in forest green), and a cluster including all *Myioborus* samples from Colombia, Ecuador, and western Venezuela including *M. o. ornatus, M. o. chrysops,* and *M. m. bairdi*).

**Figure 3.**
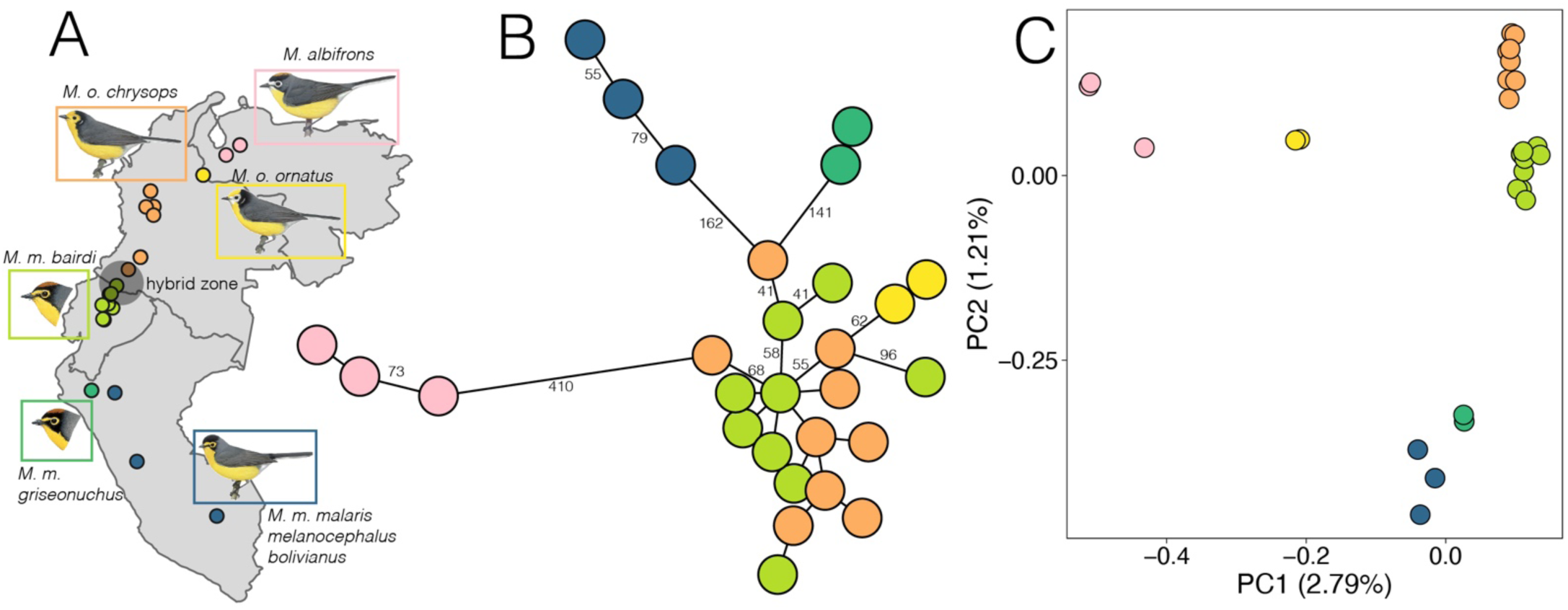
Genetic structure among the plumage groups of the *ornatus-melanocephalus* complex. **A)** Sampling localities colored by geographic groups. The approximate location of the hybrid zone is denoted as a gray circle. B) Network of mitochondrial genome haplotypes. The length of the edges connecting haplotypes are proportional to mutational steps. The specific number of mutations is shown only for edges with over 40 mutations for clarity. **C)** *angsdPCA* plot summarizing genetic variation across 12,624,354 sites.

In the genome-wide PC1/PC2 space, samples generally clustered by plumage and geography (Figure 3), although some groups with marked plumage differences appear close to each other (e.g., the hybridizing *chrysops* and*. bairdi*). Consistent with the analyses of mitochondrial genomes, *M. albifrons* appears as a clear cluster in the PC1/PC2 space, showing a strong differentiation from all the other taxa along PC1. PC2 shows a strong correlation with latitude (Supplementary Figure 2), with all southern Andes populations (southernmost *M. melanocephalus* taxa) being differentiated along this axis. Both plumage groups from the Peruvian Andes, the chestnut-crowned (*griseonuchus*), and the black-crowned (*malaris,melanocephalus, bolivianus*) taxa appear close together in the PC1/PC2 space, in contrast to their clearer separation in the mitochondrial haplotype network (Figure 3).

Additionally, in contrast to the mitochondrial haplotype network, *M. o. ornatus* does not appear to cluster with *ornatus-melanocephalus* populations occurring north of Ecuador. Instead, the *M. o. ornatus* cluster shows intermediate PC1 values between these populations and *M. albifrons*. The PC2/PC3 and PC3/PC4 spaces do not clear clustering by plumage groups but suggest some clustering of individuals collected in the same locality (Supplementary Figure 2).

### Landscape of divergence between hybridizing *Myioborus*

Out of 102 outlier 10-kb windows between these taxa, seven were not found in large regions of elevated divergence common to *Setophaga* warbler comparisons (Baiz et al. 2021). These divergent windows unique to this *Myioborus* pair fell on chromosomes 1, 1A, and 3 (Figure 4A). Comparing 10-kb windows between the hybridizing *Myioborus* species to non-hybridizing sister species *S. castanea* and *S. striata*, we found that only three windows were in the top 0.1% of *Myioborus* windows and outside the top 90% of *Setophaga* windows (Figure 4B). Two of the three were among the seven windows identified in the previous step (Figure 4C, D). The third was in a large region of elevated differentiation on the Z chromosome (Figure 4E). We note that this conservative approach focuses on the strongest outlier regions, yet more subtle differences could also exist and be relevant to shaping species differences in our focal *Myioborus* taxa.

**Figure 4.**
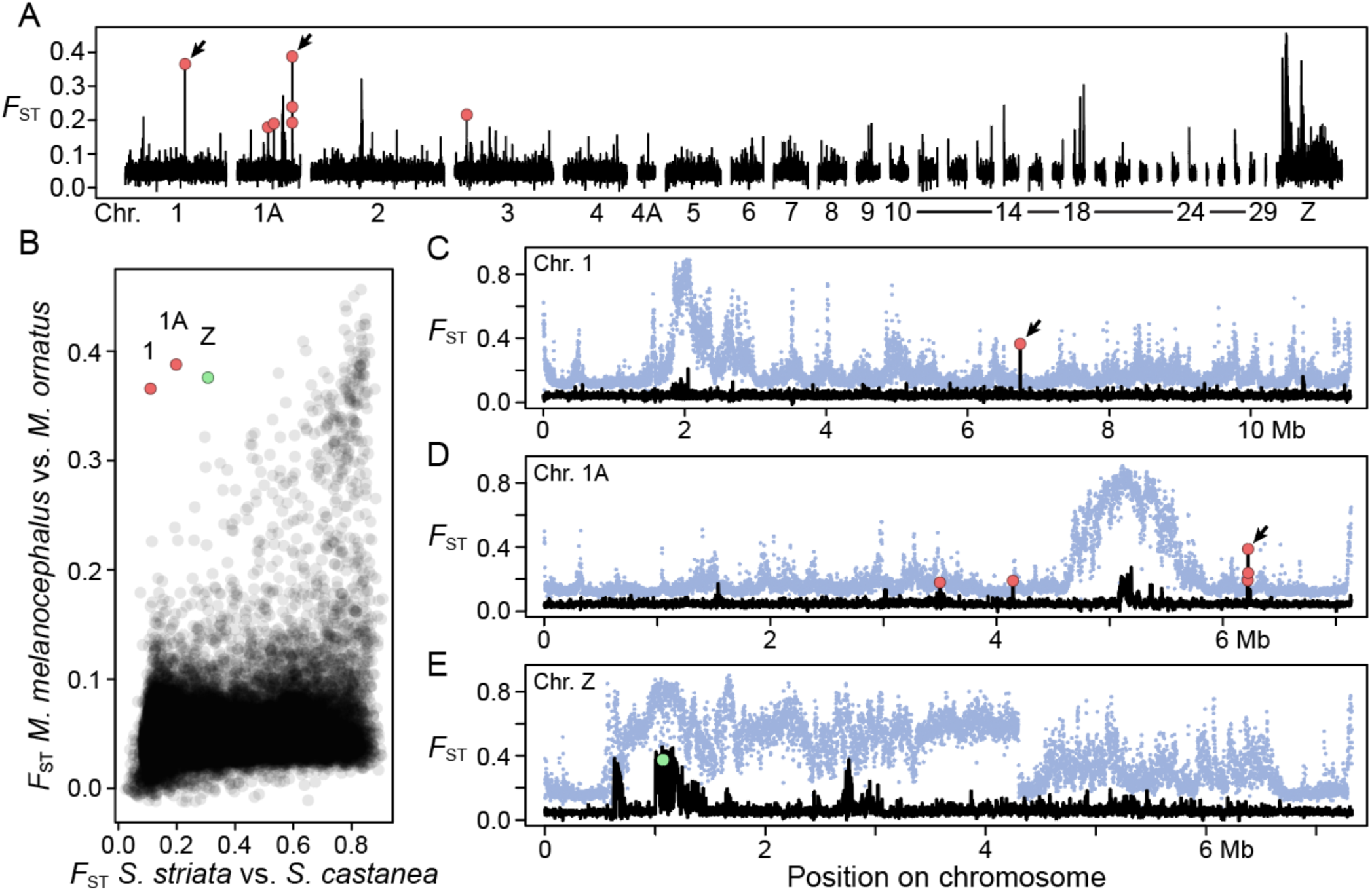
The landscape of divergence between the hybridizing *M. o. chrysops* and *M. m. bairdi* is characterized by overall low F_ST_ values with few identifiable peaks of high differentiation. At the top (**A**), the general Manhattan plot showing variation in F_ST_ values across 10 kbp windows, with the chromosome name in the x-axis. The panels (**C-E**) below show a zoom-in to regions with differentiation peaks across different chromosomes (1, 1A, and Z), including a comparison with the pattern for a *Setophaga* species pair (**B**). The F_ST_ values for the *Myioborus* pair are shown in black, and for *Setophaga* in red. This comparison is also illustrated as a scatterplot of F_ST_ values corresponding to the same windows for the *Myioborus* and *Setophaga* comparisons, highlighting the three outliers for which *Myioborus* has a peak of differentiation not seen in the *Setophaga* comparison

The chromosome 1 outlier window was more than 100 kb from the nearest annotated protein-coding gene in the yellow-rumped warbler genome. The highest chromosome 1A outlier, which also had the highest F_ST_ in the genome, partially overlapped the coiled-coil domain-containing 91 gene or CCDC91, a gene producing a protein involved in transport between the Golgi apparatus and lysosomes (Stelzer et al. 2016). The other two chromosome 1A peaks are not within 20 kb of any gene. The closest gene, approximately 30 kb away, is KITLG (KIT ligand), a protein-coding gene involved in multiple processes including cell survival and proliferation, hematopoiesis, gametogenesis and melanogenesis (Stelzer et al. 2016). The chromosome 3 peak fell within the gene C1D, which encodes for a protein that is DNA binding and that induces apoptosis (Stelzer et al. 2016).

Two regions of elevated F_ST_ in the Z-chromosome appeared to be consistent with chromosomal inversions (Figure 5A). Moving along the chromosome, F_ST_ abruptly increased and then decreased, indicating potential inversion breakpoints (Knief et al. 2024). To explore this possibility, we calculated principal components from SNP data in the putative inversions and flanking sequences (Figure 5, B-E). Consistent with inversions, the two differentiated regions showed three distinct clusters of individuals in PC space, predicted to correspond to the two inversion haplotypes plus heterozygotes (Nowling et al. 2020). In the case of the larger putative inversion, which showed the clearest clustering, PC1, which explained nearly 80% of all genetic variation among the individuals in the region, also showed a strong association with latitude (Figure 5F). *M. o. chrysops* and *M. m. bairdi* samples collected outside the hybrid zone had different predicted inversion haplotypes, based on PC1 clustering, while those collected in the hybrid zone were either predicted to be heterozygotes or have the opposite-than-expected haplotype.

**Figure 5.**
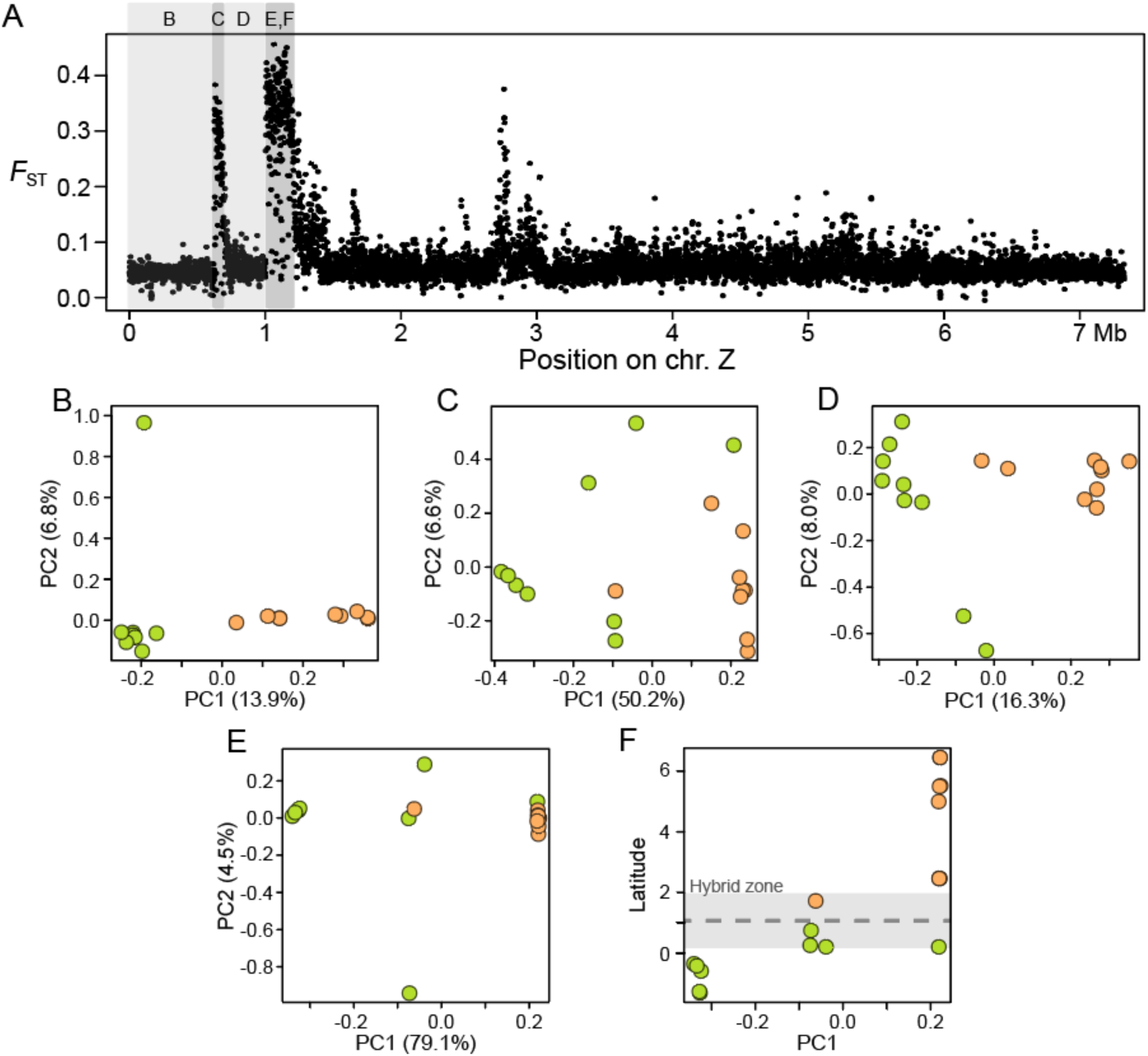
Putative inversion in the Z-chromosome **A)** Manhattan plot showing variation in FST across the Z-chromosome. The sections of the chromosome for which a PCA was performed are marked in two different shades of grey, with the letter corresponding to the other panels of the figure. In the darker shade of gray, the sections that we indetified as putative inversions (C and E,F). The panels E and F correspond to the same Z chromosome region. **B to E)** PCAs specific for each region of the chromosome Z. Dots are colored by taxa, with orange corresponding to *chrysops* and lime green to *bairdi*. Note that in C and E, there are three clearly defined clusters across the PC1 **F)** Correlation between PC1, corresponding to the same region as E, and latitude of collection of the corresponding specimens (Supplementary Table 1). The approximate location of the hybrid zone is marked in a gray shade, with the approximate center in a black dotted line (Céspedes-Arias et al. 2021)

## Discussion

We integrated phylogenetic and population-level genomic analyses to gain insights into the diversification of a tropical montane avian radiation characterized by head plumage pattern variation. This is the first genome-scale analysis of the *Myioborus*, which is the youngest genus in the rapid Parulidae radiation. Our phylogenetic study of *Myioborus* warblers, widely distributed across Neotropical mountain ranges, also sheds light more broadly on the evolutionary establishment of montane avifaunas. This genus, like warblers in general, is characterized by its high diversity of plumage patterns. By focusing on one species complex within *Myioborus* that exhibits striking geographic variation in plumage color, including two taxa with conspicuous differences in head color that hybridize, our study also contributes to our understanding of plumage diversification. We discuss our results for these different scales of analyses below.

Although topological incongruence is common in rapid radiations (Gwee et al. 2020), the tree topologies we inferred using UCEs and mitogenomes were largely concordant and agreed with previous phylogenetic hypotheses, especially at older nodes in the genus (Pérez-Emán 2005; Lovette et al. 2010; Zhao et al. 2025). Two notable common features of our tree topologies were that (1) the earliest split separated *M. pictus* (which occurs furthest north at the lowest elevation) from the rest of the genus, and (2) the next-earliest split separated *M. miniatus*, a widely distributed mid-elevation species, which was sister to an upper montane clade formed by allospecies that replace one another across different Neotropical mountain ranges, with some degree of elevational sympatry in contact zones (Jablonski et al. 2020). However, the elevational distribution of *Myioborus* species reveals a more complex pattern than a simple mid-elevation versus high-elevation dichotomy. Species from the Tepuis and Coastal Venezuela inhabit mountain tops but at lower absolute elevations than their Andean relatives due to differences in mountain height between these ranges. Also, upper elevation taxa expand their elevational ranges into mid-elevations where *M. miniatus* is absent, suggesting competitive interactions shape these distributions (Jablonski et al. 2020). These patterns support a scenario where an ancestral *Myioborus* species colonized South America via mid-elevations, followed by diversification into higher elevations in different mountain systems. Under this scenario, the widespread *M. miniatus* has maintained gene flow across its range, while upper-elevation specialists have become isolated by intervening lowlands, driving their allopatric diversification (Flantua et al. 2019). This evolutionary history, reminiscent of a taxon cycle pattern (Ricklefs and Bermingham 2002), supports deep divergence between clades that replace each other elevationally (Pérez-Emán 2005), a pattern observed in other Neotropical montane birds (Benham et al. 2015; Cadena et al. 2019).

Within the well supported high-elevation clade, we found some uncertainty in the position of *M. brunniceps* and *M. torquatus* which are, respectively, the southernmost and northernmost taxa in this group (an incongruence also inferred in a family-wide study by Zhao et al. 2025). However, given differences in support values, and extremely short internodes in this section of the mtDNA tree, we interpret our results as a whole as supporting a position of *M. torquatus* as sister to the rest of the upper montane taxa, as inferred in the UCE tree. This relationship, in turn, would suggest a north to south colonization of high elevation peaks in the Neotropical region during the diversification of these warblers, as has been inferred for other montane birds (Cadena et al. 2007; Weir et al. 2008; Gutiérrez-Pinto et al. 2017).

Within the upper montane group, two clades with clear geographic affinities are well supported across both analyses: a Pantepui plus Coastal Venezuela group (*M. pariae, M. castaneocapilla, M. cardonai, M. albifacies*) and a Tropical Andes group (*M. albifrons, M. ornatus, M. melanocephalus*). Within this first clade, we found that the Coastal Venezuela taxon is sister to the species allopatrically distributed across different tepuis, suggestive of *in situ* diversification in the Pantepui region, which is characterized by an old geological origin and high levels of endemism (Borges et al. 2018). Within both the Pantepui and Coastal Venezuela group and the Tropical Andes, we found topological incongruence between UCEs and mitogenomes. Several factors could cause this discordance (Toews and Brelsford 2012), including past introgression of mitochondrial or nuclear DNA (Toews et al. 2013; Kearns et al. 2018; Berbel-Filho et al. 2022) or a high degree of incomplete lineage sorting (DeRaad et al. 2023). Both explanations are plausible, especially for a rapidly diversifying group. The phylogenomic analyses presented herein do not allow us to distinguish these possibilities.

All three of our phylogenies strongly support the existence of a Tropical Andes clade formed by *M. albrifrons*, and the geographically variable *M. melanocephalus* and *M. ornatus*. Overall, we found that each geographically circumscribed plumage group appears as a cluster in our mitogenomes network. A notable exception is the *M. ornatus* and *M. melanocephalus bairdi* group, which show marked differences in plumage but show low divergence in mitochondrial DNA (also see Céspedes-Arias et al. 2021). This contrasts with the PCA patterns summarizing variation across the nuclear genome, in which *M. o. ornatus* appears as a distinct cluster, in an intermediate position along the PC1 between the *bairdi-chrysops* group and *M. albifrons*.

Notably, in our genus-wide phylogenomic analyses, we found that the phylogenetic affinities of *M. o. ornatus* differ between our inferences based on mitochondrial and nuclear data. Given the short internodes in this section of the trees, suggestive of rapid diversification, incomplete lineage sorting is a plausible explanation for this apparent mito-nuclear discordance (Degnan and Rosenberg 2009; Meleshko et al. 2021; DeRaad et al. 2023). Another, not exclusive possibility, is that historical introgression between some of these *Myioborus* lineages may be also contributing to observed differences in patterns of variation in mitochondrial and nuclear DNA. A more extensive geographic sampling of whole-genome data within these groups, along with targeted analyses to detect introgression, will be necessary to disentangle these possibilities.

Our analyses at three different evolutionary scales support that the hybridizing *M. m. bairdi* and *M. o. chrysops* are sister taxa and show overall low genetic differentiation, corroborating previous work focused on few mitochondrial markers (Pérez-Emán 2005; Céspedes-Arias et al. 2021). These two, likely young, taxa are characterized by clear differences in head coloration, mirroring the genus level pattern of plumage variation.

In describing the landscape of divergence between these two taxa, we found that the levels of differentiation across the genome are highly heterogeneous, with some well-identified peaks. These peaks, for the most part, fall within large regions of divergence shared with pairs of *Setophaga* warblers (Baiz et al. 2021). As Baiz et al. (2021), we interpret this shared differentiation across independent pairs as a function of a common genomic architecture, reduced recombination, and linked selection (Cruickshank and Hahn 2014; Burri et al. 2015).

Genomic regions of reduced recombination like centromeres have previously been shown to associate with increased differentiation between diverging taxa because linkage is much greater, which amplifies the effects of divergent selection (Burri et al. 2015; Van Doren et al. 2017). In the case of very closely related diverging taxa like *M. melanocephalus* and *M. ornatus*, these peaks are fairly narrow, which may provide an opportunity to identify genes underlying linked divergent selection. For instance, there are only four genes in the center of the putative centromeric peak on chromosome 2: SLC9A3, BRD9, COL15A1, and TPPP.

In the *Myioborus* landscape of divergence we also identified several peaks that are not shared with the *Setophaga*, and that could represent regions underlying the phenotypic differences between these young hybridizing taxa, most notably, head coloration. One of these regions, located on chromosome 1A (Figure 4, blue arrow), contains the gene CCDC91, which codes for a protein whose function is not well understood, but that has been linked to acrokeratoelastoidosis, which is a rare skin condition in humans (Zhu et al. 2024). This gene has previously been found in regions of high differentiation between species with melanin-based plumage differences, including munias (*Lonchura*) that differ in head and body color (Stryjewski and Sorenson 2017) and wagtails (*Motacilla*) that differ in head color (Semenov et al. 2021). In the case of the wagtails, the two subspecies differ primarily in the extent of melanin around their eyes and in their cheek patches, which are the same patches that differs between *chrysops* and *bairdi*. Thus, as with other melanogenesis genes that have been found to be commonly diverged in independent avian pairs (e.g. MC1R, ASIP) (Toews et al. 2016; Campagna et al. 2017, 2022; Baiz et al. 2021), CCDC91 may have been independently recruited for similar pigmentation phenotypes in both *Motacilla* and *Myioborus*.

We found that the Z-chromosome shows a higher overall differentiation relative to other autosomes, a pattern prevalent among bird species pairs (Irwin 2018), and particularly marked across hybrid zones (Hooper et al. 2019) in which Z-linked loci often show reduced introgression (Carling and Brumfield 2008; Taylor et al. 2014; Walsh et al. 2016; Ottenburghs 2022). This pattern of higher differentiation relative to autosomes is also observed in the X chromosome in organisms with an XY sex determination system (Presgraves 2018). In general, the pervasiveness of this pattern is consistent with an important role of sex chromosomes in speciation (Irwin 2018; Presgraves 2018). Moreover, in the presence of gene flow, these patterns could be interpreted as evidence that sex chromosomes are less prone to introgression across species boundaries. Cline analyses in hybrid zones across multiple bird taxa appear to support this idea, as Z-linked loci often show reduced introgression relative to autosomes (Carling and Brumfield 2008; Taylor et al. 2014; Walsh et al. 2016; Hooper et al. 2019; Ottenburghs 2022), and furthermore, some suggest a direct involvement of Z-linked loci in genetic incompatibilities (Lopez et al. 2021).

Given the existence of a hybrid zone between *chrysops* and *bairdi*, it is possible that gene flow could be contributing to the different levels of divergence in the Z-chromosome relative to autosomes. However, the observed pattern alone does not allow us to draw this conclusion because this could also be a product of demography or linked selection within populations (Presgraves 2018).

Besides having overall higher differentiation in the *chrysops-bairdi* pair, the Z-chromosome contains some of the most prominent divergence peaks, which could be caused by chromosomal inversions (Figure 5). Specifically, we noted a sharp drop of F_ST_ at the end of the divergent regions (i.e., a table-top shape), which together with the region-specific PCA patterns in which three discrete clusters are identifiable, is consistent with regions of suppressed recombination (Nowling et al. 2020). Chromosomal inversions, because of their effect drastically reducing recombination, can play an important role restricting gene flow across hybrid zones (Hooper et al. 2019), and contribute to speciation (Hoffmann and Rieseberg 2008). The hybrid zone between *M. m. bairdi* and *M. o. chrysops* is narrow elevationally but latitutidanally extensive, stretching across more than 200 km. Within this zone, intermediate phenotypes are polymorphic and predominant, though hybridization remains restricted to a fraction of the range of these taxa (Céspedes-Arias et al. 2021, Céspedes-Arias, Campagna *et al*, preprint), suggesting that hybridization is quite recent or that the width of the hybrid zone is being constrained by selection (Barton 1979; Gay et al. 2008). Because it falls on the Z chromosome, the putative inversion we identified may play an outsized roll in contributing to reduced fitness in hybrids (Hooper et al. 2019), and therefore partial reproductive isolation between these taxa. Some next steps to evaluate the potential role of these inversions in hybridization dynamics are to corroborate their existence using complimentary evidence, e.g., long-read sequencing, and to explore fitness consequences of the inversion haplotypes and heterozygotes.

We used whole-genome resequencing data for most species in the genus *Myioborus* to gain insights about the evolutionary history of a radiation of resident Neotropical warblers at different time scales. Our work also sets the stage for future avenues of research. First, a question that arises is what the role of introgression is driving phylogenomic discordance. Second, our work suggests that the two hybridizing taxa withing these genus (*M. m. bairdi* and *M. o. chrysops*), have shallow divergence in both their mitochondrial and nuclear genome and the landscape of divergence between them led us to identify one candidate gene for their differences in melanin-based plumage patches. Future studies should obtain whole genome data for hybrids with a diversity of plumage phenotypes to evaluate the correlation between genetic variation in this locus and the presence/absence of specific patches, as well as to identify other candidate genes, by leveraging the wide variation that exists in this hybrid zone. Lastly, we identified a putative inversion in the Z-chromosome that could be impacting hybridization dynamics. By using whole genome data from individuals collected across the hybrid zone it will be possible to assess whether there is evidence of reduced introgression of the Z-chromosome, and gathering long read data could corroborate the existence of the hypothesized inversion.

## Author contributions

LNCA, KB, LC and DT conceived the study; LNCA and AMC conducted fieldwork to collect the *bairdi* and *chrysops* samples, with logistical support from CDC and EB. AW, KB and LNCA conducted lab work. KB conducted the bioinformatic processing of raw data and data analyses. KB, LNCA and DT produced figures. LNCA led the writing of the original draft with major contributions from KB and DT, and additional input from all other authors. All authors contributed to the interpretation of results and to reviewing and editing the manuscript.

## Acknowledgements

We are thankful to museum collection that loaned the tissues for whole genome re-sequencing: AMNH, FMNH, KU, LSUMZ, MSB, UWMB, and YPM. The following people provided valuable support during fieldwork to collect the samples for *bairdi* and *chrysops* in Colombia and Ecuador used here: D. Ocampo, M.A. Meneses, P. Pulgarín, A. Mendoza, and F. González. We are also thankful to many local guides, landowners, nature reserve officials and other members of local communities who supported our work in Colombia and Ecuador. The Universidad de los Andes (Colombia) and Universidad Tecnológica Indoamérica (Ecuador) provided essential support to obtain export permits for the *chrysops* ad *bairdi* tissues. We thank Bronwyn Butcher for her invaluable technical support during lab work in the Fuller Lab (Cornell Lab of Ornithology). A portion of the computations for this research were performed on Pennsylvania State University’s Institute for Computational and Data Sciences’ Roar Collab supercomputer. We also thank John M. Bates for his valuable suggestions during the preparation of this manuscript.

## Data availability

Raw sequence data is available from NCBI (BioProjects PRJNA630247 and X). All other data and code necessary to reproduce the analyses from this study are available from Dryad (DOI: 10.5061/dryad.j3tx95xt).

---

**Supplementary Figure 1:**
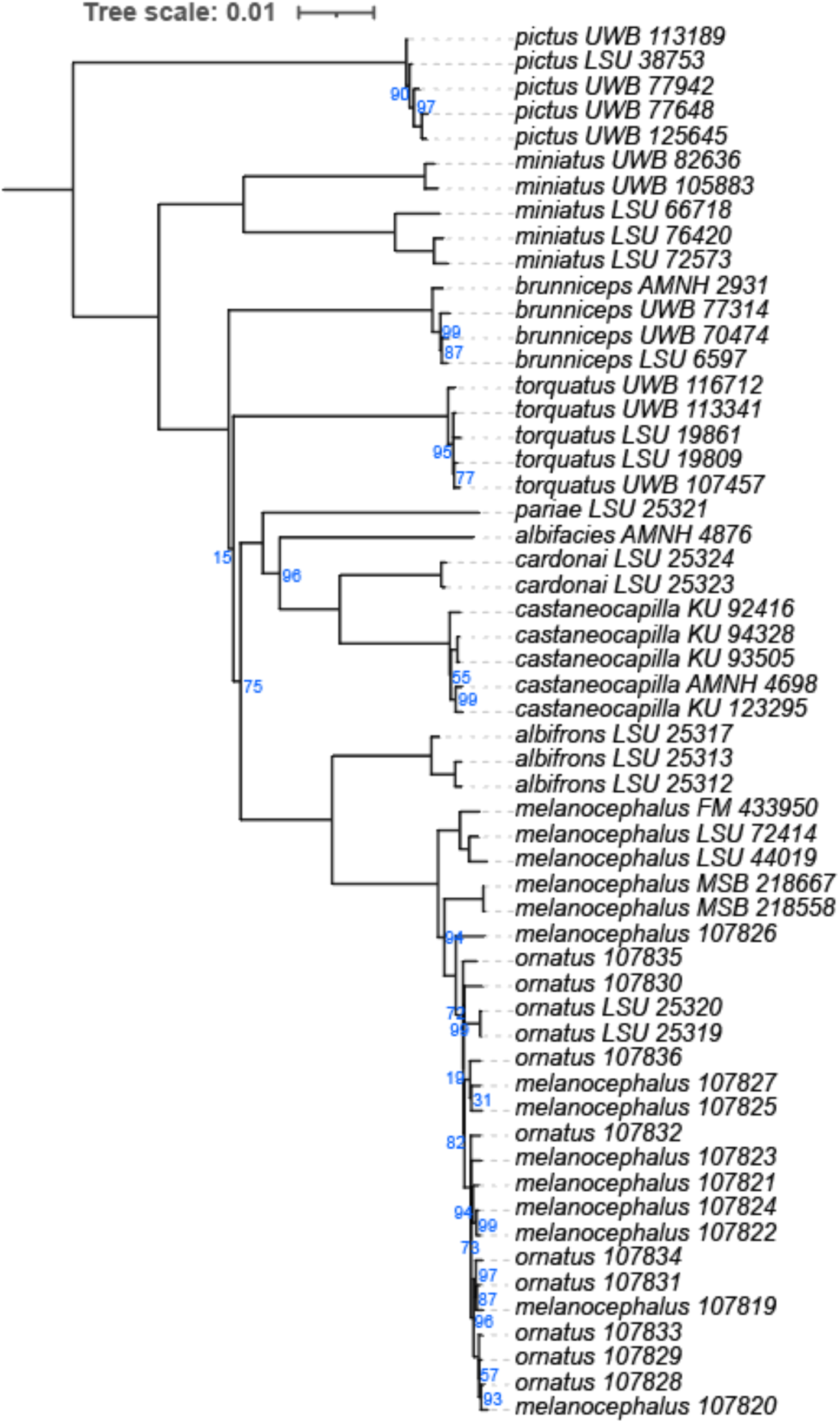
Mitogenomes *Myioborus* phylogeny with branch support values. Only values below 90 are shown. The tip labels correspond to individuals as named in the Supplementary Table 1.

**Supplementary Figure 2:**
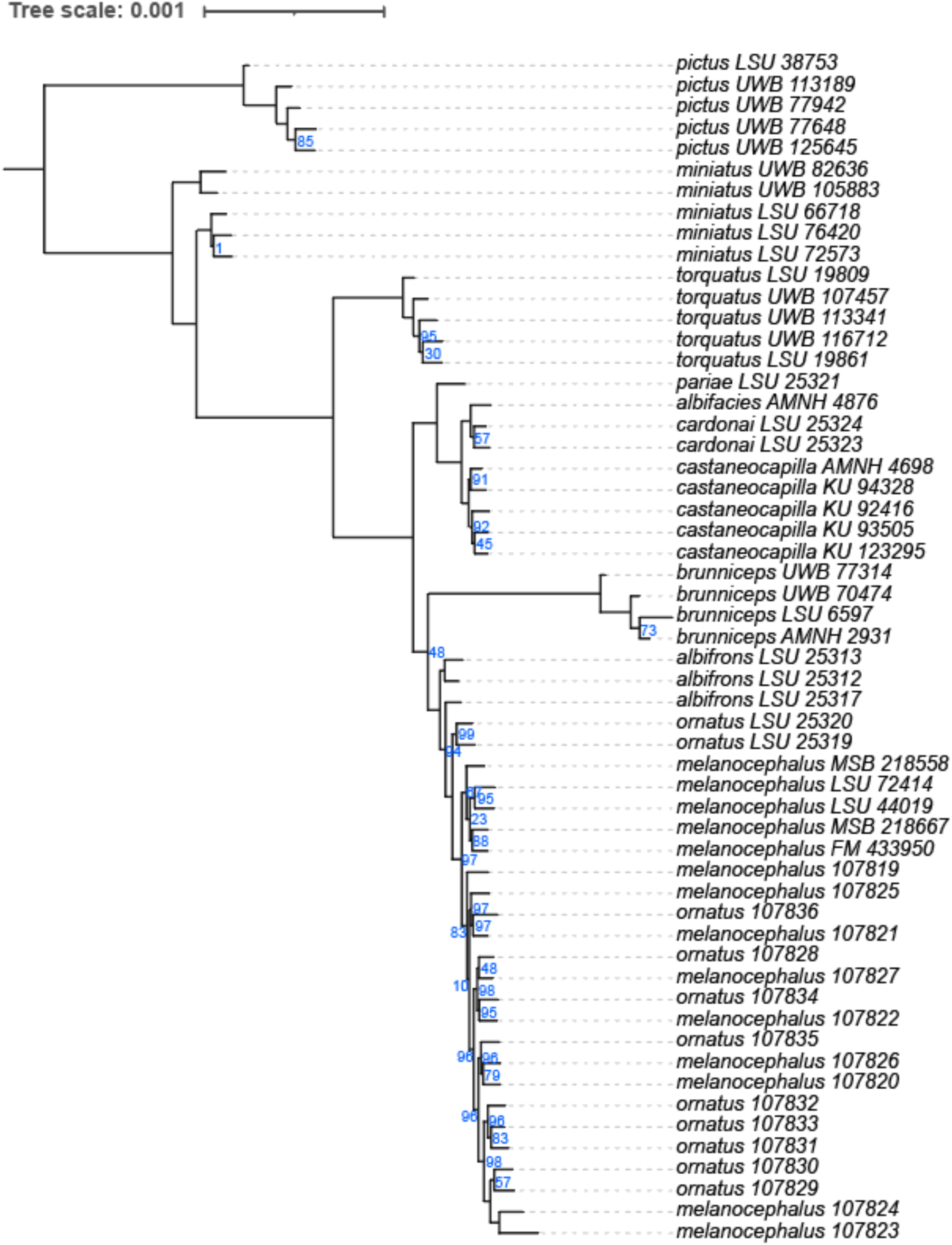
Concatenated UCEs *Myioborus* phylogeny with branch support values. Only values below 90 are shown. The tip labels correspond to individuals as named in the Supplementary Table 1.

**Supplementary Figure 3.**
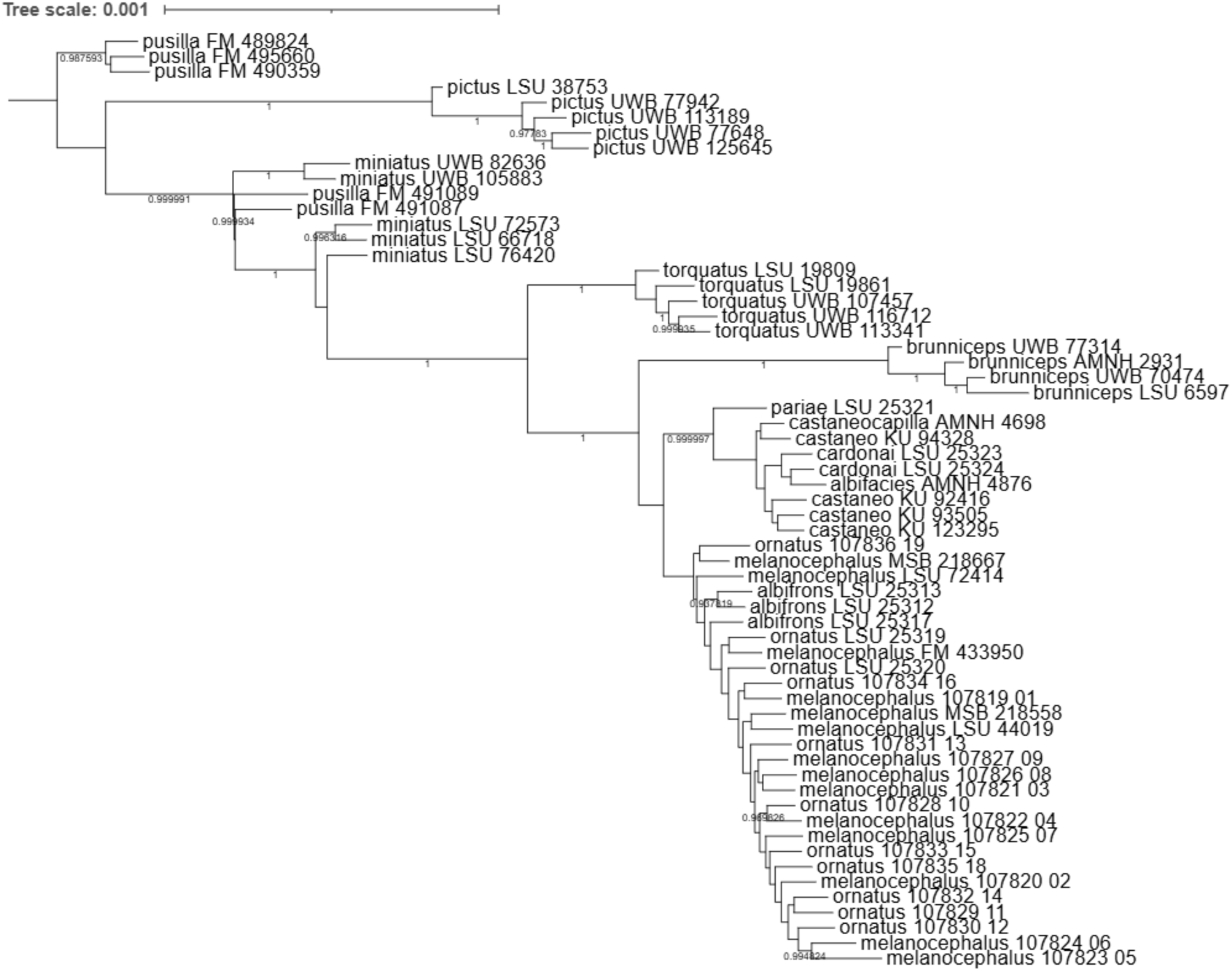
UCE species tree of *Myioborus*, infered using ASTRAL-III. Branch upport values below 90 are shown, and tip labels correspond to individuals as named in the Supplementary Table 1.

**Supplementary Figure 3.**
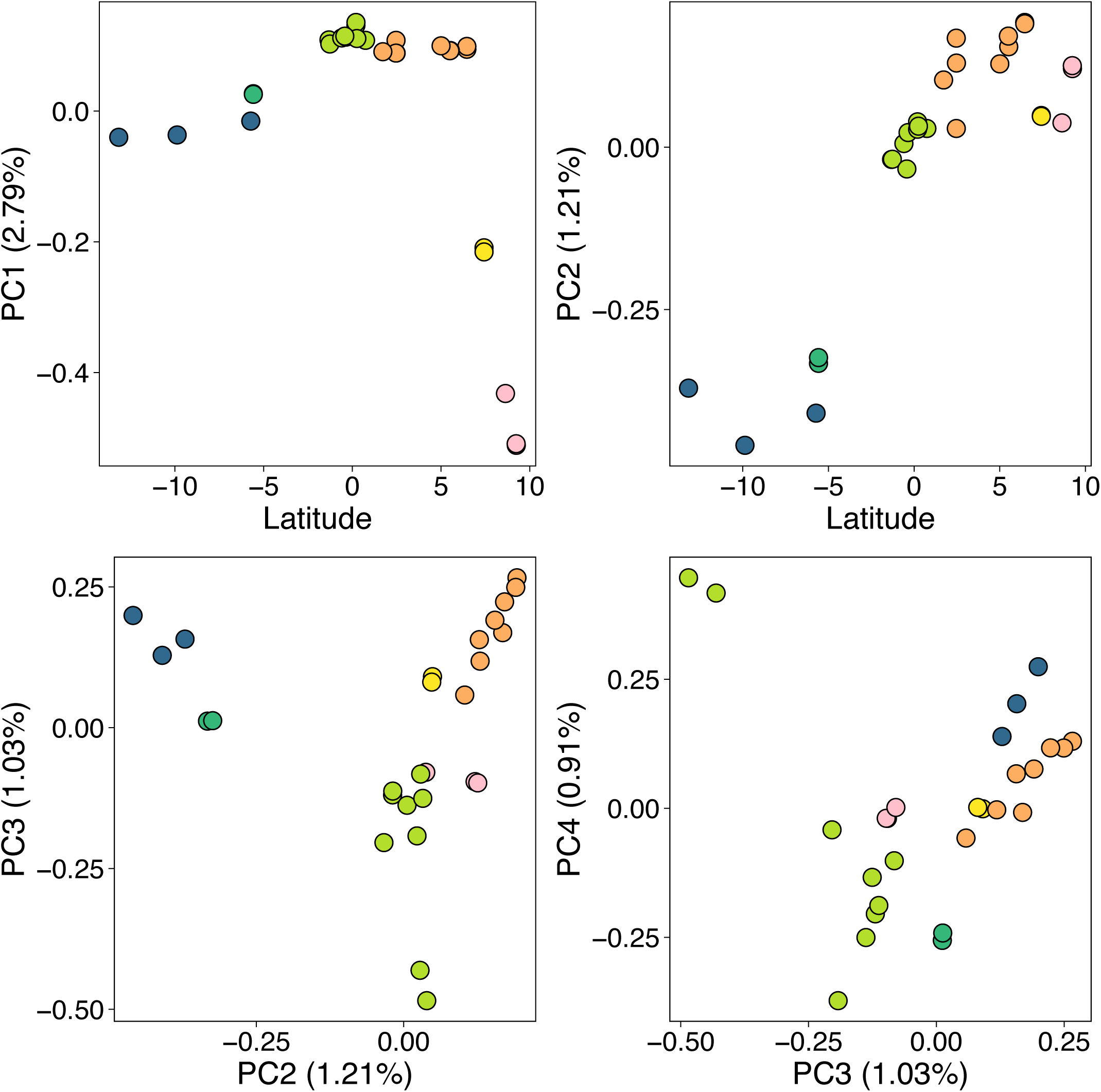
Genetic structure within the group encompassing *M. ornatus, M. melanocephalus* and *M. albifroms* complex as summarized by a PCA. Top panels: relationship between the two first principal components and latitude. PC2 shows a strong latitudinal pattern.

